# A state-dependent neural circuit resolves approach–avoidance conflicts

**DOI:** 10.64898/2026.02.05.704045

**Authors:** Devika Bodas, Şevval Demirci, Marine Balcou, Lisa Scheunemann, Carolina Rezaval

## Abstract

To survive, animals must balance opportunity and risk, yet how internal motivational state biases this trade-off remains poorly understood. Here we show that hunger gates nociceptive avoidance in *Drosophila* to enable food approach under conflict. Using a behavioural assay in which flies must cross an electric shock barrier to reach food, we find that starvation promotes shock tolerance only when food-related olfactory cues are present. This state- and context-dependent behavioural shift is mediated by a defined neuromodulatory circuit. Leucokinin neurons encode hunger and convey this information to PV5K1 lateral horn output neurons, which integrate internal state signals with food odour cues and suppress shock-responsive FB2B neurons in the ventral fan-shaped body. Accordingly, electric shock–evoked activity in FB2B neurons is suppressed in starved flies exposed to food cues. These findings identify a circuit mechanism by which motivational state gates aversive processing to bias action selection towards goal-directed behaviour.

**Teaser:** Hunger enables flies to suppress nociceptive avoidance when food cues signal reward availability via a defined neuromodulatory circuit.

## Introduction

The ability to resolve conflicts between competing drives, such as pursuing rewards while avoiding harm, is fundamental for survival. Reward-predictive cues promote approach behaviours driven by motivation, whereas threat cues elicit defensive responses that minimise risk. In natural environments, however, these signals often co-occur, generating approach–avoidance conflicts in which organisms must integrate internal state with external sensory information to prioritise action. How the brain performs this integration to flexibly bias behaviour without compromising survival remains a central question in neuroscience.

Foraging provides a clear example of this trade-off. Seeking food frequently exposes animals to danger and aversive sensory cues, yet avoiding harm can result in the loss of essential resources. Although defensive behaviours typically dominate behavioural hierarchies, they can be suppressed or gated when motivational drive or expected reward value outweighs perceived risk (*1*, *2*). Across species, animals will tolerate pain, threat, or uncertainty to secure food, mates, or shelter, particularly when resources are scarce (*3–5*). Hunger is a strong driver of such reweighting processes (*6–8*), biasing behaviour towards goal pursuit even at a cost (*5*, *9–14*). In rodents, hunger modulates inflammatory pain responses (*15*, *16*) and in humans, it attenuates pain responses (*17*). These observations point to a general principle: internal state can reshape how aversive sensory information is evaluated to favour goal pursuit. Yet how exactly motivational signals override nociceptive circuits, and how they bias behavioural choice without globally suppressing defensive responses, remains unclear. The fruit fly *Drosophila* provides an ideal system for dissecting these mechanisms at single-cell and circuit-level resolution(*18*, *19*), while exhibiting conserved neural principles of action selection shared with mammals (*1*, *20*). In flies, electric shock evokes robust innate avoidance (*21*, *22*) and is processed by defined central brain circuits, including neurons of the ventral fan-shaped body (vFB), a structure implicated in behavioural selection and state-dependent control (*22–25*). Hunger and satiety states are regulated by multiple neuropeptidergic systems, including leucokinin (LK), which modulates feeding, sleep, and thermal sensitivity (*26–29*). However, how hunger-dependent signals interface with nociceptive avoidance circuits to bias behavioural choice under conflict has remained unresolved.

Here, we show that hunger biases approach–avoidance decisions through a defined neuromodulatory pathway linking metabolic state to sensory integration. We identify LK neurons as a key source of hunger signals acting upstream of a specific class of lateral horn output neurons (LHONs), PV5K1, which integrate internal state with food-related sensory cues and inhibit shock-responsive neurons in the vFB. Through this pathway, hunger selectively attenuates nociceptive avoidance when food cues are present, enabling flies to tolerate noxious stimuli during reward pursuit. Hunger thus shifts cost–benefit weighting by suppressing immediate avoidance only under conditions in which goal attainment is possible.

## Results

### Hunger promotes food pursuit despite nociceptive cues

We developed an approach–avoidance paradigm in which flies must choose between avoiding an electric shock (ES) zone or crossing it to access food (Fig. 1A). The yeast-enriched food source emitted salient olfactory cues detectable before commitment, creating a conflict between nociceptive avoidance and reward pursuit.

**Figure 1:**
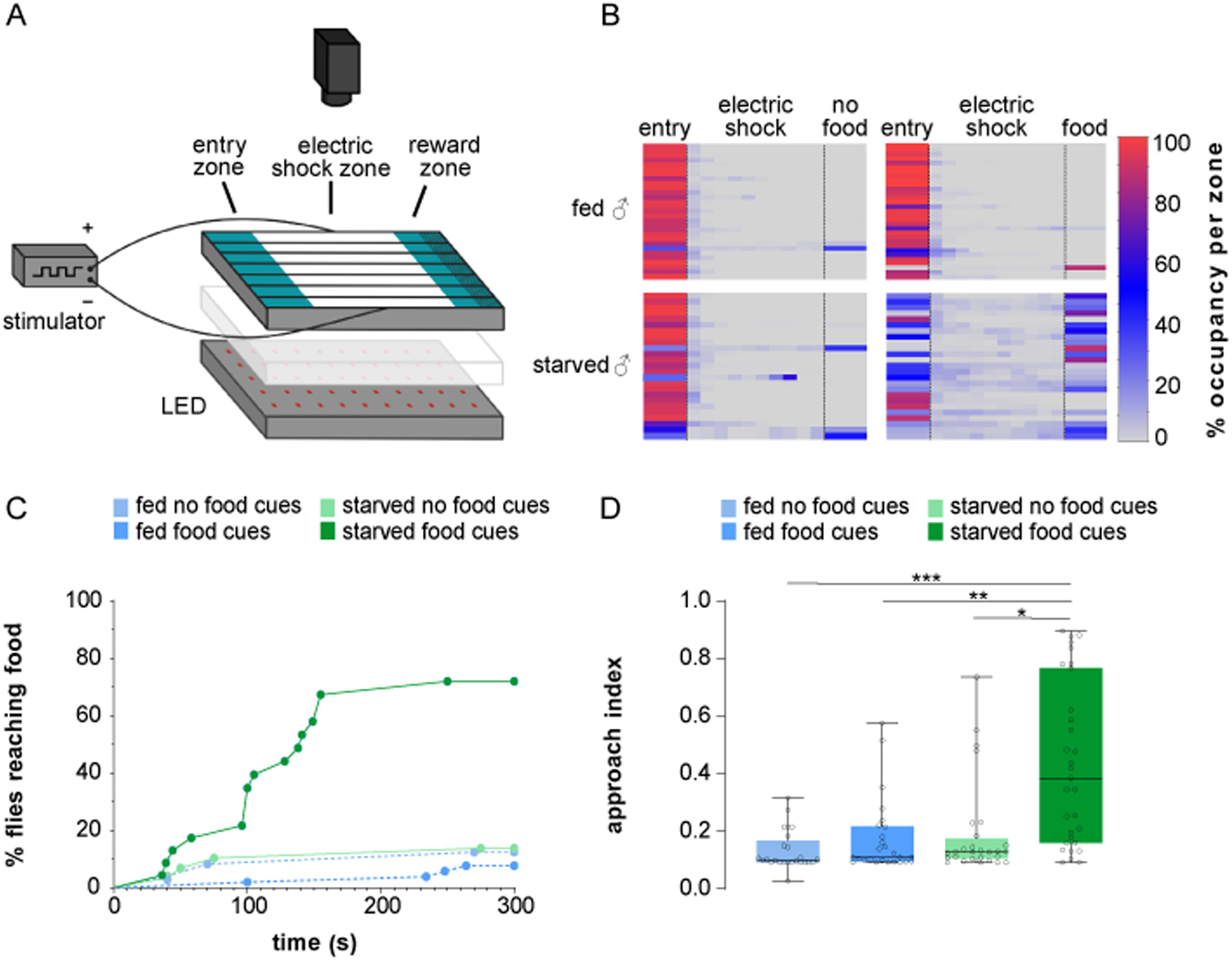
Starvation enhances food approach despite nociceptive cues. **(A)** Schematic of a novel approach–avoidance assay designed to probe decision-making under conflict in *Drosophila*. Individual flies traverse an arena comprising an entry zone, an aversive electric shock zone, and a reward zone containing yeast-based food. Flies can choose to avoid the electric shock by remaining in the entry zone or cross the aversive shock zone to obtain food. **(B)** Heatmaps depict the spatial occupancy of individual flies across the arena over the 5-min trial, with colour intensity representing the percentage of time spent in each location. In the presence of food cues, 24-h-starved male flies spent more time in the electric shock and reward zones compared with fed controls and starved flies tested without food in the reward zone, reflecting an increased propensity to traverse aversive terrain to access reward. All data were acquired under 75□V electric shock (n = 28-29 flies per condition). **(C)** Temporal dynamics of reward approach during exposure to 75□V electric shock, shown as the cumulative percentage of flies reaching the reward zone over time. Fed males are indicated by dotted lines (dark blue, food cues present; light blue, food cues absent), whereas 24-h-starved males are indicated by solid lines (dark green, food cues present; light green, food cues absent). Starvation increased the proportion of flies reaching the reward zone when food cues were available, whereas fed flies and starved flies tested without food cues showed minimal reward approach (n = 28-29 flies per condition). **(D)** Quantification of reward approach using an approach index during 75□V electric shock exposure. Fed males (dark blue, food present; light blue, food absent) and 24-h-starved males (dark green, food present; light green, food absent) are shown. The approach index was significantly elevated in starved flies when food cues were available, whereas all other conditions exhibited comparably low approach under aversive stimulation (n = 28–29 flies per condition). Box plots indicate the median and interquartile range; whiskers denote the minimum and maximum values, and individual data points represent single flies. Statistical significance was assessed using Kruskal–Wallis tests followed by Dunn’s post hoc comparisons (*□p□<□0.05, **□p□<□0.001, ***□p□<□0.0001).

We quantified food approach across ES intensities using a binomial generalized linear model incorporating shock voltage, feeding state, and food cues. Increasing ES intensity reduced the probability of reaching the reward zone across all conditions (Suppl. Fig. 1). However, when food cues were available, starved flies were more likely to approach the reward zone despite high ES intensities. At 75□V, fed flies and starved flies tested without food cues showed strong avoidance, whereas starved flies presented with food cues continued to approach the reward zone; this condition was therefore used for all subsequent experiments.

Detailed behavioural analyses revealed that at 75□V, only a small fraction of fed flies reached the reward zone during the 5-min assay, whereas ∼70% of 24-h-starved flies crossed the ES zone and reached the reward, typically within the first 2.5□min (Fig. 1B–C; Suppl. Videos 1, 2). In the absence of food odours, starved flies resembled fed controls, avoiding the ES zone and failing to approach the reward. Consistently, an approach index based on spatial occupancy increased with starvation only when food cues were present, with no difference between fed and starved flies tested without food cues (Fig. 1D; see Methods).

Together, these results show that hunger enhances motivational drive rather than reducing shock sensitivity, enabling suppression of nociceptive avoidance only when reward-predictive sensory cues signal the availability of food cues.

### FB2B neurons in the ventral fan-shaped body encode electric shock and mediate innate avoidance

Previous work has shown that vFB neurons are activated by electric shock and are required for innate nociceptive avoidance (*22*), although the contribution of defined cell types remains unresolved. Using behavioural assays and functional imaging, we identified FB2B neurons (Fig. 2A), a defined layer 2 vFB population, as a substrate for innate ES avoidance. Silencing FB2B neurons increased the approach index in both fed and starved flies, irrespective of availability of food cues (Fig. 2B–D). Baseline locomotor speed was unaffected by FB2B silencing (Suppl. Fig. 2), excluding altered motor output as an explanation for the behavioural effects.

**Figure 2.**
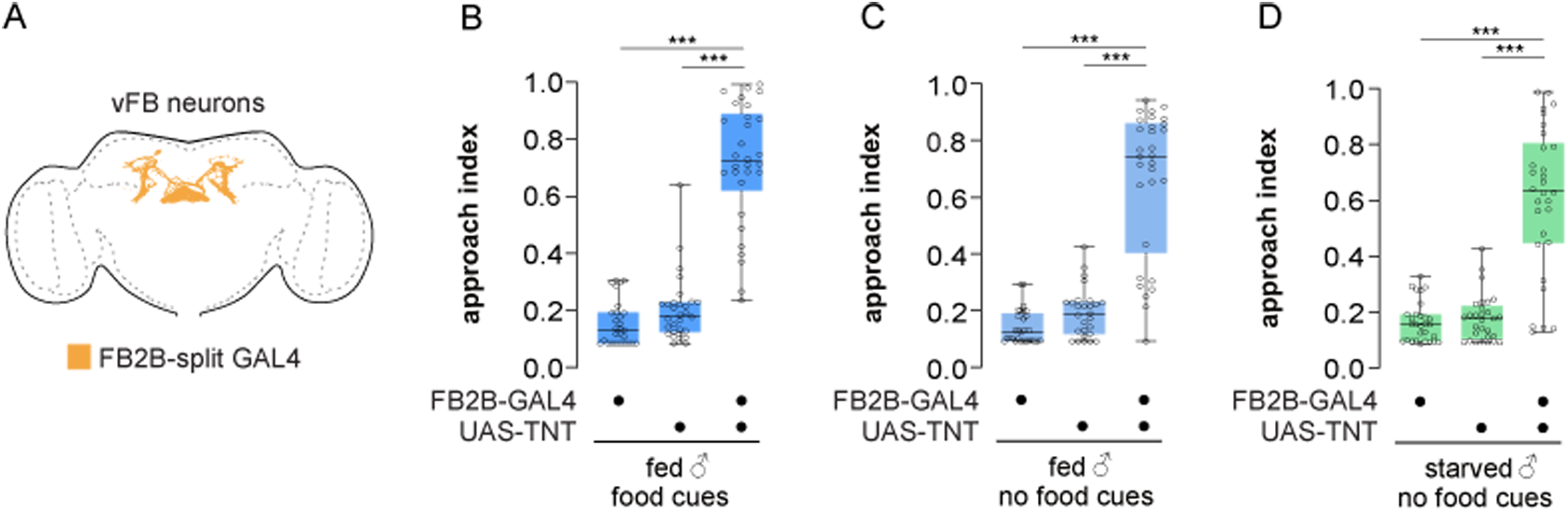
A subset of vFB neurons is required for starvation-driven reward approach under nociceptive threat. **(A)** Schematic showing expression of the FB2B-split GAL4 driver labelling FB2B ventral fan-shaped body (vFB) neurons. **(B–D)** Blocking synaptic transmission in FB2B neurons using tetanus toxin light chain (TNT) increases reward approach under aversive conditions. Approach indices were quantified in FB2B-GAL4 > UAS–TNT male flies and their respective genetic controls tested during 75□V electric shock exposure under fed conditions in the presence (B, dark blue) or absence (C, light blue) of food cues, and under 24-h starvation in the absence of food cues (D, light green). In all conditions, FB2B silencing elevated approach compared to genetic controls. Box plots show the median and interquartile range; whiskers indicate minimum and maximum values; points represent individual flies. Statistical significance was assessed using Kruskal–Wallis tests followed by Dunn’s post hoc comparisons (*p□<□0.05, **p□<□0.001, ***p□<□0.0001). n = 27–30 flies per condition.

Consistent with a role in nociceptive processing, functional calcium imaging of FB2B neurons expressing GCaMP7b revealed robust activation in response to ES in fed flies (Fig. 5C). Together, these results demonstrate that FB2B neurons encode electric shock and are required for innate ES avoidance.

### Hunger-dependent food approach despite nociceptive cues is mediated by leucokinin neurons

Because hunger promotes food approach despite nociceptive cues (Fig. 1), we sought the neural signal that conveys internal metabolic state to the circuits governing this decision. Leucokinin (LK) neurons encode hunger state and modulate hunger-dependent behaviours in *Drosophila* (*26–30*). We therefore tested whether LK neurons promote food approach by attenuating nociceptive avoidance during reward pursuit.

Silencing LK neurons using LK-GAL4–driven Kir2.1 significantly reduced food approach in starved flies (Fig. 3A-B), without affecting basal locomotion (Suppl. Fig 3A-B). Consistent with a role for LK signalling itself, RNAi-mediated knockdown of LK similarly attenuated food approach despite electric shock (Fig. 3C). To test sufficiency, we thermogenetically activated LK neurons using dTrpA1, a manipulation well suited for engaging neuropeptide-dependent modulation (*31*). Acute activation of LK neurons increased food approach in fed flies at 30°C but not 24°C (Suppl. Fig. 3D) but only in the presence of food-related sensory cues (Fig. 3D–E), without affecting walking velocity (Suppl. Fig. 3C).

**Fig 3:**
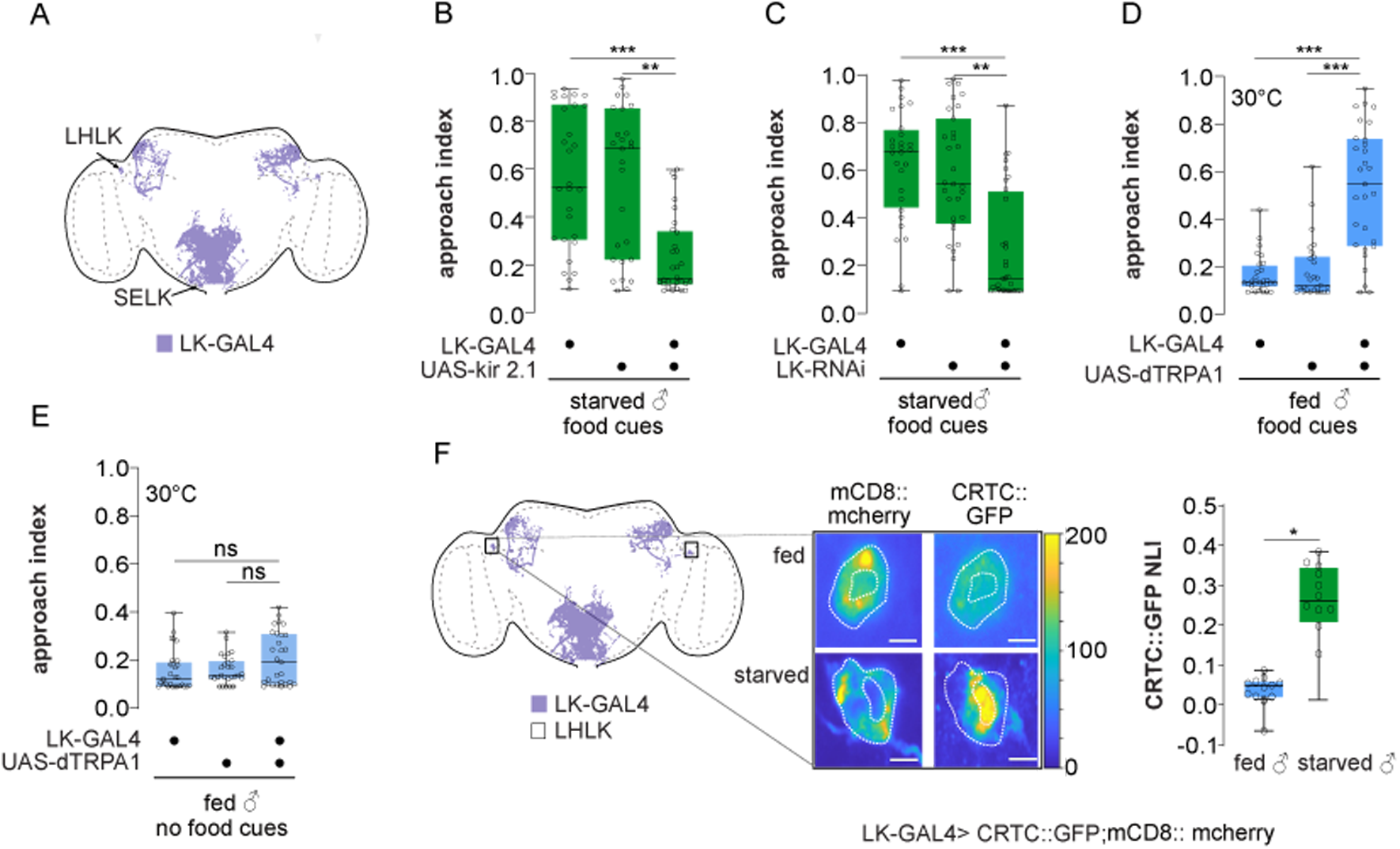
Leucokinin signalling favours hunger-driven food approach over avoidance. **(A)** Expression pattern of LK-GAL4, labelling leucokinin (LK) neurons in the lateral horn (LHLK) and the subesophageal zone (SELK). **(B)** Silencing LK neurons with the inwardly rectifying potassium channel Kir2.1 reduces hunger-driven food approach. Approach indices were measured in 24-h-starved LK-GAL4 > UAS–Kir2.1 males and their respective genetic controls tested in the presence of food cues (green) during 75□V electric shock exposure (n = 27–30 flies per condition). **(C)** Downregulation of leucokinin (LK) expression in LK neurons using RNAi suppresses hunger-driven food approach. Approach indices were measured in 24-h-starved LK-GAL4 > UAS–LK RNAi males and their respective genetic controls tested in the presence of food during 75□V electric shock exposure (n = 28–32 flies per condition). **(D)** Thermogenetic activation of LK neurons promotes food approach in fed flies. Approach indices were measured in fed LK-GAL4 > UAS–dTRPA1 males and their respective genetic controls tested at 30°C in the presence of food cues during 75□V electric shock exposure (n = 27–29 flies per condition). **(E)** Thermogenetic activation of LK neurons does not promote approach in the absence of food cues. Approach indices were measured in fed LK-GAL4 > UAS–dTRPA1 males and their respective genetic controls tested at 30°C without food cues (light blue) during 75□V electric shock exposure (n = 28–29 flies per condition). **(F)** LHLK neurons show increased activity under starvation. Neural activity was assessed in LHLK neurons (schematic, left) using the transcriptional reporter CRTC::GFP. Membranes were labelled with mCD8::mCherry to delineate nuclear and cytoplasmic compartments (white dotted outlines). Representative images illustrate distinct CRTC localisation patterns in fed and starved flies, with corresponding nuclear localisation indices (NLI) shown alongside; heat maps indicate GFP pixel intensity (1–200 a.u.). Quantification of CRTC::GFP nuclear localisation is shown on the right. Starvation significantly increased CRTC::GFP nuclear localisation in LHLK neurons, consistent with elevated neuronal activity under nutrient deprivation (n = 8 flies per condition). Box plots indicate the median and interquartile range; whiskers represent the minimum and maximum values; points represent individual flies. Statistical significance was assessed using Kruskal–Wallis tests followed by Dunn’s post hoc comparisons for panels B–E and a Mann–Whitney test for panel F (right) (^∗^p < 0.05; ^∗∗^p< 0.001; ^∗∗∗^p< 0.0001).

Using the activity-dependent CRTC::GFP reporter (*32*), we observed a robust starvation-induced increase in activity in a lateral horn LK population (LHLK neurons; Fig. 3F) but not in a suboesophageal zone LK population (SELK neurons; Suppl. Fig. 3E-F) targeted by the LK-Gal4. The feeding state dependent changes in activity of LHLK neurons is consistent with prior work (*28*, *30*). Because LK-GAL4 also labels both clusters of neurons LHLK along with, future intersectional approaches will be required to assign causality specifically to LHLK neurons.

Together, these findings identify LK neurons as a key source of hunger-state signals that bias behavioural choice, enabling food pursuit to override nociceptive avoidance when reward-predictive cues are available.

### Leucokinin signalling via LHON neurons mediates hunger-dependent food approach despite aversive cues

Because LK neurons encode hunger state and are embedded within the lateral horn (*28*, *30*, *33*), we reasoned that hunger-dependent biasing of food approach during sensory conflict is likely implemented at the level of lateral horn output neurons (LHONs). The lateral horn transforms ethologically relevant sensory cues into behaviour, e.g. navigation towards attractive odours, positioning its outputs to bias downstream action selection circuits (*23*, *34*, *35*).

PV5K1 neurons project from the lateral horn to layer 2 of the vFB, which contains shock-responsive FB2B neurons, and no other characterised LHON population innervates this layer (Fig. 4A; (*35*)). PV5K1 neurons thus emerge as a candidate pathway through which hunger-related signals could reduce shock avoidance during food approach.

**Figure 4:**
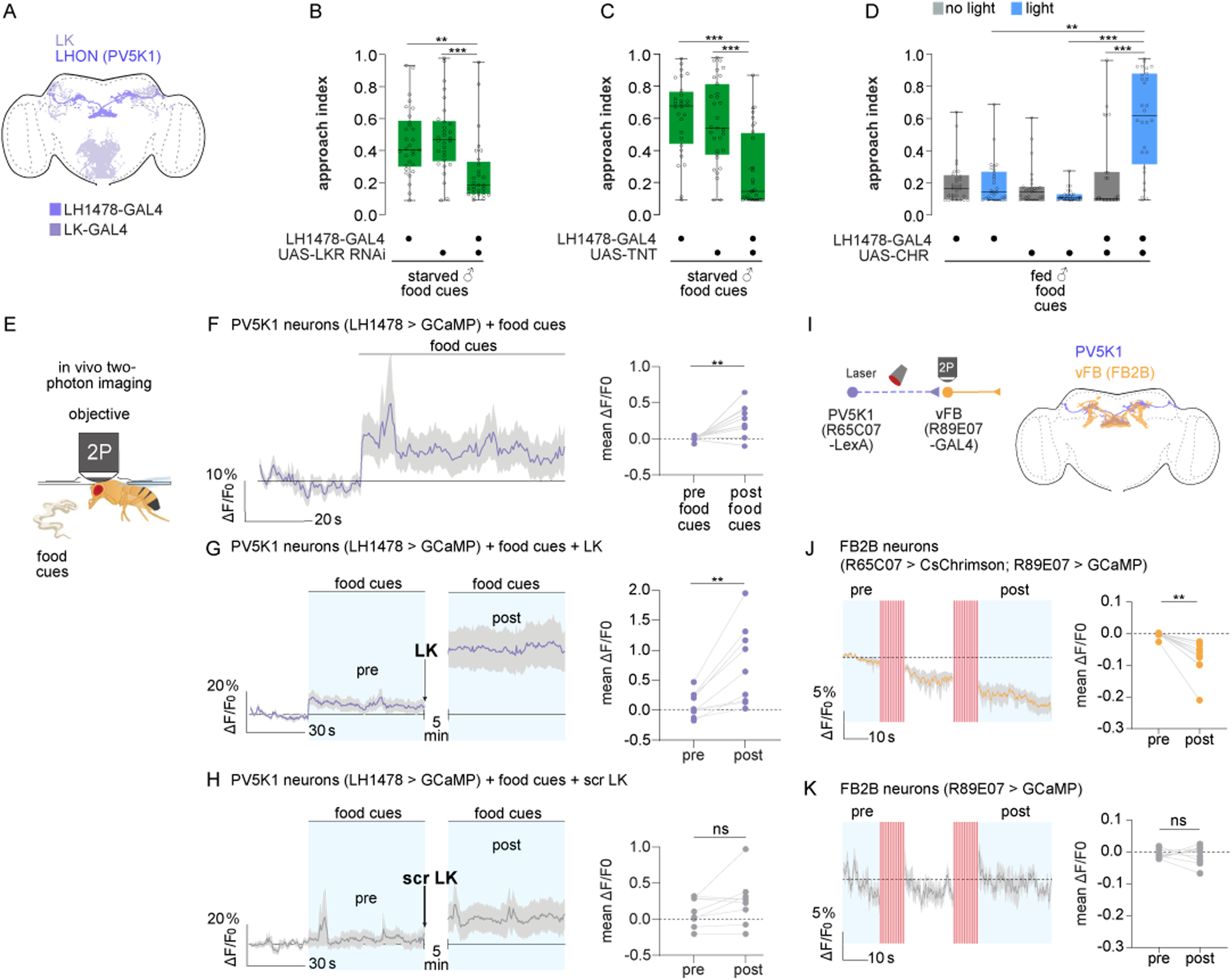
A leucokinin-responsive lateral horn circuit promotes food approach despite nociceptive cues. **(A)** Expression patterns of leucokinin (LK) neurons (lavender) and PV5K1 lateral horn output neurons labelled by LH1478-GAL4 (violet) in the brain. **(B)** LK signalling in PV5K1 neurons is required for starvation-driven reward approach despite nociceptive cues. Approach indices were measured in 24-h-starved male flies and their respective genetic controls tested in the presence of food cues during 75□V electric shock exposure following knockdown of the leucokinin receptor (LH1478-GAL4 > UAS–LKR RNAi) (n = 29–30 flies per condition). **(C)** PV5K1 neurons are required for starvation-driven reward approach despite nociceptive cues. Approach indices were measured in 24-h-starved male flies and their respective genetic controls tested in the presence of food cues during 75□V electric shock exposure following synaptic silencing of PV5K1 neurons (LH1478-GAL4 > UAS–TNT) (n = 29–30 flies per condition). **(D)** Optogenetic activation of PV5K1 neurons promotes food approach over avoidance in fed flies. Fed males expressing CsChrimson in PV5K1 neurons (LH1478-GAL4 > UAS–Chr) and their respective genetic controls were tested in the presence of food cues during 75□V ES exposure, with or without light stimulation (n = 26–29 flies per condition). Boxes indicate the median and interquartile range; wwhiskers represent the minimum and maximum values; points represent individual flies. Kruskal-Wallis tests followed by Dunn’s post hoc comparisons were performed for panels B-D (^∗^p < 0.05; ^∗∗^p< 0.001; ^∗∗∗^p< 0.0001). **(E)** Schematic of the *in vivo* calcium imaging setup used to monitor neural activity during food odour presentation. **(F-H)** Left, Δ*F*/*F*_0_ calcium traces from PV5K1 neurons (LH1478-GAL4>UAS-GCaMP6f) in fed males in the presence of food cues (**F**), before and after application of 100 µM LK peptide (**G**), or 100 µM control scrambled LK peptide (**H**). Responses in PV5K1 neurons to food cues increase upon LK application. The sample sizes represent biologically independent flies. Solid lines and shaded areas of live-imaging traces indicate mean□±□S.E.M, respectively. Right, mean Δ*F*/*F*_0_ values during pre-stimulus and post-stimulus periods (quantification windows indicated as blue boxes, n = 9 - 10 flies per condition). Paired t-test was performed in F. Paired Wilcoxon signed-rank test was performed in G, H. (^∗^p < 0.05; ^∗∗^p < 0.001; ^∗∗∗^p < 0.0001). **(I)** Schematic illustrating the strategy used to assess functional connectivity between PV5K1 neurons and vFB (layer 2, FB2B) neurons (left). Brain schematic showing the anatomical overlap of PV5K1 neurons (violet) and vFB neurons (yellow) (right). **(J-K)** Left, ΔF/F_0_ calcium traces from vFB neurons (layer 2, FB2B) (R89E07 > GCaMP6f) in males with (**J**) or(R89E07 > GCaMP7b) in males without (**K**) optogenetic activation of PV5K1 neurons using CsChrimson (red bars). Activation of PV5K1 neurons results in a reduction of vFB calcium signals in FB2B neurons consistent with an inhibitory functional interaction. Traces show mean ± s.e.m. Right, quantification of mean ΔF/F_0_ values during the pre-stimulation window and following the second stimulation (quantification windows indicated as blue boxes, n = 12 for J, n = 10 for K). Statistical significance was assessed using a paired t-test for J and a Mann–Whitney test for K (*p□<□0.05, **p□<□0.001, ***p<□0.0001).

Consistent with this hypothesis, knockdown of the leucokinin receptor (LKR) in PV5K1 neurons significantly impaired food approach in starved flies (Fig. 4B). Similarly, silencing PV5K1 neurons reduced approach behaviour under starvation (Fig. 4C), whereas acute optogenetic activation of PV5K1 neurons was sufficient to promote food approach in fed flies despite electric shock (Fig. 4D). These manipulations did not affect baseline locomotor speed (Suppl. Fig. 4A-D), indicating a specific effect on behavioural choice. By contrast, knockdown of LKR in FB2B neurons had no effect on approach–avoidance behaviour (Suppl. Fig. 4E-F), arguing against direct leucokinin modulation of shock-responsive neurons.

Given the role of LHONs in processing innate olfactory cues (*36*, *37*), we next asked whether PV5K1 neurons integrate hunger signals with food-related sensory information.

Live calcium imaging revealed that PV5K1 neurons were activated by yeast odours in fed flies (Fig. 4E–F). Bath application of leucokinin, used to mimic starvation, significantly potentiated these odour-evoked responses (Fig. 4G), whereas scrambled control peptides had no effect (Fig. 4H). Importantly, LK application in the absence of food odours was insufficient to elicit PV5K1 activity (Suppl. Fig. 4G). Together, these results indicate that LK signalling does not directly drive PV5K1 activity but instead enhances the gain of food-evoked sensory responses under hunger.

To determine whether PV5K1 neurons directly influence nociceptive circuits, we optogenetically activated PV5K1 neurons while monitoring FB2B activity (Fig. 4I). PV5K1 activation suppressed calcium activity in FB2B neurons (Fig. 4J-K), revealing an inhibitory functional connection between these populations.

Together, these findings indicate that PV5K1 lateral horn output neurons integrate leucokinin-dependent hunger signals with food odours to suppress nociceptive avoidance and favour food seeking.

### Hunger gates nociceptive avoidance through context-dependent suppression of FB2B

Our findings indicate that flies override nociceptive avoidance only when hunger coincides with food-related sensory cues (Fig. 1), consistent with modulation of aversive processing via the LK–PV5K1–vFB pathway.

To test whether hunger-dependent gating occurs at the level of FB2B neurons, we examined whether their responses to electric shock are modulated in a context-dependent manner. Because vFB layer 2 neurons have been shown to exhibit shock-evoked calcium responses (*22*), we performed *in vivo* calcium imaging of FB2B neurons while delivering ES under the microscope, in the presence or absence of yeast-derived food cues (Fig. 5A-B). FB2B neurons were activated by electric shock, showing an increase in GCaMP signal during the shock window relative to baseline (Fig. 5C). In the absence of food cues, ES-evoked FB2B responses were similar in fed and starved flies (Fig. 5D, F). In contrast, when food cues were present, ES-evoked FB2B activity was strongly suppressed in starved flies (Fig. 5E-F), matching the observed behaviour (Fig. 1).

**Figure 5:**
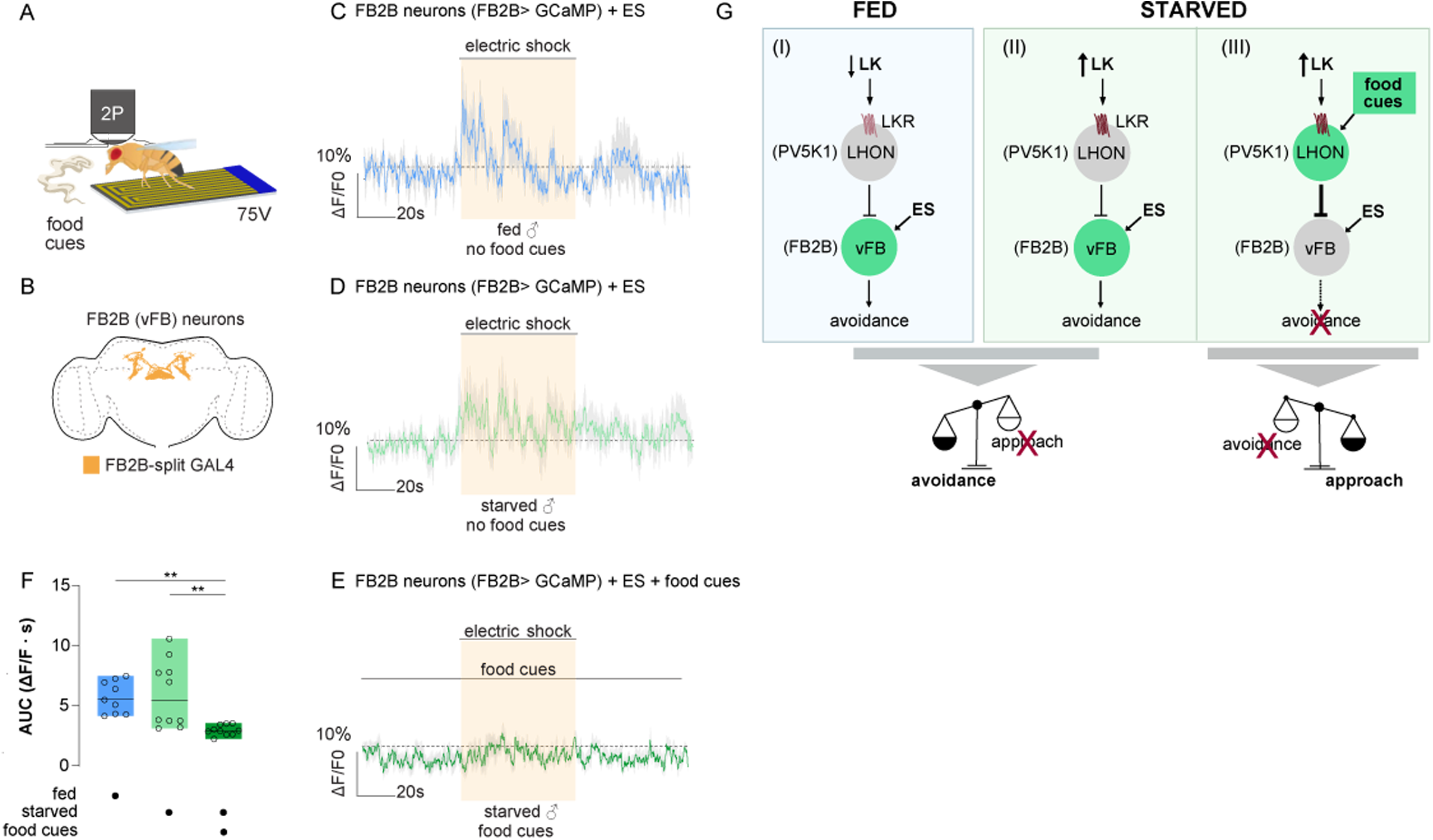
A lateral horn-fan shaped body pathway enables hungry flies to overcome aversive cues to reach food. **(A)** Schematic of the *in vivo* two-photon imaging set up used to monitor neural responses during electric shock delivery in behaving flies. **(B)** Expression pattern of FB2B-split GAL4 labelling FB2B vFB neurons.**(C–E)** ΔF/F_0_ average traces of FB2B□>□GCaMP7b signals in fed flies (C) and in starved flies tested in the absence (D) or presence (E) of food cues, shown before, during, and after electric shock presentation (yellow shaded area). FB2B neurons exhibit robust shock-evoked calcium responses in fed flies and in starved flies tested without food cues, whereas shock responses are strongly suppressed in starved flies when food cues are present. Solid lines and shaded areas indicate mean□±□s.e.m., respectively (n= 9-10 flies per condition). **(F)** Quantification of vFB neural activity expressed as area under the curve (AUC) of ΔF/F_0_ responses during shock exposure. vFB response.s are similar between starved and fed flies without food but are selectively reduced by availability of food cues in starved flies. Box plots show the median response for each condition (n = 9–10 flies per condition). One-way ANOVA test followed by Tukey’s HSD post hoc comparisons was performed (^∗^p < 0.05; ^∗∗^p < 0.001; ^∗∗∗^p < 0.0001). **(G)** Proposed model for state- and food-dependent gating of nociceptive avoidance by the LK–LHON–vFB pathway. In fed flies (I), LK signalling to lateral horn output neurons (LHONs; PV5K1) is reduced, resulting in low PV5K1 activity (grey). Ventral fan-shaped body FB2B (vFB) neurons remain active during electric shock (ES), driving avoidance behaviour. In starved flies without food cues (II), LK signalling is elevated but insufficient to activate PV5K1 neurons. As a result, FB2B activity and shock avoidance are largely preserved. In starved flies exposed to food cues (III), enhanced LK signalling together with sensory food input strongly activates PV5K1 neurons (green). This increased PV5K1 activity suppresses FB2B (vFB) responses to ES, attenuating nociceptive avoidance and biasing behaviour toward food approach despite aversive stimulation. Grey indicates neural inhibition; dark green indicates neural activation.

Altogether, our results identify a hunger-dependent neuromodulatory circuit in which LK–LKR signalling enhances food-cue responsiveness in PV5K1 neurons, leading to hunger-specific suppression of nociceptive avoidance via FB2B inhibition (Fig. 5G).

## Discussion

Animals must face conflicts in which pursuing reward carries risk, requiring neural systems that can flexibly prioritise behaviour without globally suppressing defensive responses. Here, we identify a circuit mechanism by which internal metabolic state selectively biases action selection during such conflicts in *Drosophila.* Hunger does not blunt nociceptive processing; instead, it gates avoidance in a context-dependent manner, permitting tolerance of aversive stimuli only when reward-predictive cues are available.

Using a novel approach–avoidance assay, we show that starvation promotes food approach across noxious electric shock exclusively in the presence of food odours. In their absence, starved flies behave indistinguishably from fed controls, favouring nociceptive avoidance. This dissociation demonstrates that hunger amplifies motivational drive rather than diminishing sensory sensitivity to aversive input. Comparable state-dependent biases in valuation and action selection, without loss of threat detection, have been described across species (*6*, *38–40*).

In *Drosophila*, several neural populations spanning sensory and central brain circuits mediate electric shock responses ^(*22*, *41–44*)^. Our findings identify FB2B neurons in the ventral fan-shaped body (vFB) as a key node for resolving conflicts between nociceptive avoidance and reward pursuit. FB2B neurons are activated by electric shock and are required for avoidance regardless of feeding state or availability of food cues, positioning them as a sensory–motor pathway for defensive behaviour. Importantly, FB2B activity is not globally suppressed by starvation; instead, shock-evoked responses are attenuated only when hunger coincides with food cues. This context dependence mirrors behaviour and suggests that FB2B neurons act as a gate rather than a simple relay of nociceptive information.

Our data identify LK signalling as a critical source of hunger-state information that biases this gating. Previous work has established LK neurons as sensors of nutritional state and regulators of hunger-dependent behaviours, including feeding, sleep, and nociceptive sensitivity (*26–30*). In particular, lateral horn LK neurons increase activity during starvation (*30*) and are suppressed by glucose, directly encoding metabolic state in mated females (*28*). Our findings extend this framework by showing that LK signalling is required for hunger-driven food approach during approach–avoidance conflicts in males. Given that hunger in *Drosophila* is encoded by multiple neuromodulatory systems rather than a single pathway (*45*), it is likely that additional hunger signals converge on FB circuits to modulate aversive decision-making under distinct motivational contexts.

We further demonstrate that hunger-state signals are routed through a specific class of lateral horn output neurons, PV5K1(*35*). The lateral horn has emerged as a hub for transforming innate sensory cues into behavioural commands (*23*, *34*, *35*), yet how internal state information interfaces with its outputs remains unclear. PV5K1 neurons are uniquely positioned to fulfil this role: they receive lateral horn input and project selectively to layer 2 of the vFB, where FB2B neurons reside(*46*). Functional manipulations show that PV5K1 neurons are required for hunger-dependent food approach and sufficient to override avoidance in fed flies. Optogenetic and calcium imaging experiments further reveal that these neurons integrate food odours with LK signals to inhibit FB2B activity. Together, these results delineate a hunger-dependent neuromodulatory circuit in which LK–LKR signalling enhances food-cue responsiveness in PV5K1 neurons, leading to context-specific inhibition of FB2B activity (Fig. 5G). This mechanism allows starvation to gate nociceptive electric shock avoidance only when reward pursuit is possible, thereby promoting food seeking while preserving defensive responses in non-rewarding contexts. Given that vFB neurons also mediate avoidance of thermal threats (*22*), FB2B neurons likely contribute to a broader aversive control pathway.

Consistent with central hub models of action selection (*47*, *48*), our findings identify FB2B neurons as a control node that integrates nociceptive input with state- and cue-dependent modulatory signals to regulate behavioural choice. It would be interesting to test whether distinct dorsal fan-shaped body neurons that gate feeding initiation based on nutritional state and resource density (*49*)interact with FB2B avoidance circuits to coordinate risk–reward decisions across behavioural conflicts. This view aligns with accumulating evidence implicating the fan-shaped body in action selection and behavioural flexibility (*23–25*, *50*). The fan-shaped body is a core component of the arthropod central complex, a structure proposed to implement an action-selection architecture that shares organisational principles with the vertebrate basal ganglia (*51*).

Our data further show that a subset of flies crossed the shock zone at intermediate intensities even when satiated and in the absence of food cues, suggesting that avoidance is not solely governed by metabolic state. In mammals, exploratory drive biases decisions toward information seeking under uncertainty and risk (*52*, *53*), and humans will tolerate electric shocks to satisfy curiosity, engaging striatal circuitry that overlaps with hunger-driven risk taking (*54*). These findings raise the possibility that exploratory drive, like hunger, converges on fan-shaped body circuits to bias behavioural choice.

While this study focused on fly males, our findings naturally extend to questions of sex and reproductive state. Nociceptive sensitivity and risk tolerance are sexually dimorphic across species ^(*55–61*)^, often reflecting differences in motivational priorities rather than sensory detection. In *Drosophila* females, mating increases feeding drive to support egg production ^(*18*, *62–64*)^, a process known to engage leucokinin signalling (*65*, *66*). Given that we identify LK-dependent suppression of avoidance, it will be important to test whether nociceptive gating is enhanced or reconfigured in mated females, where nutritional demand is elevated. Such comparisons will determine whether the circuit logic described here represents a shared decision framework with sex-specific thresholds, or distinct implementations tuned to reproductive state.

More broadly, this work illustrates how neuromodulators bias behaviour by acting on intermediate decision nodes rather than sensory or motor endpoints. Similar architectures have been described in mammals, where hunger, stress, or motivational state modulate threat processing through defined hypothalamic and cortical pathways without abolishing nociceptive encoding (*6*, *39*, *67*). The simplicity and genetic accessibility of the *Drosophila* system allowed us to resolve this principle at single-cell resolution, providing a tractable framework for dissecting how internal state shapes adaptive behaviour under conflict.

## Materials and Methods

### Fly husbandry and strains

Flies were raised on standard diet of cornmeal/agar food in a 12-hour light-dark cycle at 25°C, or at 30°C for LK RNAi experiments. Wild type flies were all from the Canton-S (CS) strain. Flies were collected and sorted under CO2 anaesthesia post eclosion and housed in same-sex groups of 10-15. All strains used and their sources are highlighted in Suppl. Table 1. Brain expression patterns of Gal4 and Split-Gal4 lines were schematised based on FlyLight database (Rubin, G. M., et al. Cell Reports 2013).

### Trans-retinal food preparation

Trans-retinal (R2500-100MG, CAS number: 116-31-4, Sigma-Aldrich) was diluted to 50 mM in ethanol and stored at −20°C. Aliquots were wrapped in foil to protect them from light. 20 µl of 50 mM all-trans retinal factor was added to surface of food vials and allowed to dry for a day before use. All vials were wrapped with aluminium foil to prevent light exposure.

### Behavioural assays

#### Approach-avoidance behavioural assay

Five to ten days old male flies were used for all behavioural assays. Behaviour was recorded for 5 min at 60 frames per second using a Mako U-130B camera equipped with an infrared filter (BP735-40.5, Midopt). The transparent behavioural arena was positioned on top of an LED board emitting IR light. All trials were conducted between ZT1 and ZT5 to minimize circadian effects, and trial order was randomized across groups.

The approach-avoidance behavioural assay (Fig. 1A) consists of seven arenas each 12 cm long and 4 mm wide. Each arena is subdivided into an entry zone, a shock zone, and a reward zone containing high yeast food. The surface of the shock zone comprises a transparent shock grid made of a 75 mm x 50 mm polyethylene terephthalate (PET) sheet coated with a conductive indium tin oxide (ITO) 175 μm film, with a surface resistivity of 80 Ohms/Sq (Diamond Coatings Ltd., UK). A grid was laser-etched onto the ITO film to insulate the positive and negative electrodes (the lanes in the grid were 0.5 mm spaced by 0.3 mm apart). The conductive grid connected to an electrical stimulator that delivered a 1 s electric shocks every 2 s (0.5 Hz) of 75□V (unless mentioned otherwise) upon contact. The stimulator output was verified by measuring the current going through the electric shock grid using an oscilloscope, prior to doing behavioural experiments. Flies were introduced into the arena through a loading area on the left of the entry zone, separated from the main arena by a movable gate. When the flies quickly entered the entry zone, the gate was closed. The flies could then move along the arena through the shock zone up until the reward zone on the right.

To prevent the flies from avoiding the shock grid by walking on the arena lid, the underside of the lid was treated with the Sigmacote siliconizing reagent (Sigma-Aldrich, SL2), rendering the surface hydrophobic. Arenas were rinsed with water between trials and allowed to aerate for 5 min between experiments to eliminate residual olfactory cues.

Trials in which flies avoided the shocks by walking on the lid or remained immobile in the entry zone for the duration of recording, were excluded from analysis. All experiments were performed in darkness in a temperature-controlled room maintained at 25°C and 50–60% relative humidity.

#### Basal velocity assay

Flies were introduced into the approach–avoidance behavioural arena and allowed to acclimatise for 2 min. Locomotor activity was quantified by measuring fly velocity for 5 mins using EthoVision XT 18 software (Noldus Information Technology). During the assay, no electric shock was applied.

#### Starvation Protocol

Five to ten days old male flies were wet starved in 1% agar vials with filter paper soaked in water for 24 hours. Agar vials were watered 1 hour before behavioural experiments to prevent dehydration. All flies were flipped the day before starvation onto a fresh vial of food.

#### Optogenetic assay

Male flies were collected after eclosion and placed on standard cornmeal/agar food for a period of 2 days. Flies were then transferred to food containing *all*-trans retinal factor at 3 days of age. They were flipped onto fresh retinal food the day before behavioural testing. In starvation experiments, starvation agar vials were supplemented with 20 µl of 50 mM all-trans retinal.

Flies were tested in the approach avoidance behavioural setup and illuminated from underneath with 660 nm (red) light in the absence or presence of the food. They were presented with 60 Hz red light throughout the behavioural recording.

#### Thermogenetic assay

Male flies were collected after eclosion and placed on standard cornmeal/agar food at 25°C. Flies were then transferred to 30°C for 2 hours before the experiments. Flies were tested in the approach avoidance behavioural setup at 28°C in the absence or presence of the food.

### Behavioural analysis

The position of each fly in the arena was tracked throughout the experiment, and the cumulative time spent in each zone (%) were computed using the EthoVision XT18 software.

#### Approach index calculation

We developed an approach index formula to quantify approach-avoidance behaviour. To calculate the approach index (AI), the electric shock (ES) zone was divided into 10 equal subzones in Ethovision XT18. Each was assigned a weight, increasing from left (0.1) to right (1.0), to reflect the relative proximity to the reward zone. The reward zone was assigned the maximum weight of 1.1. The approach index was calculated as a weighted average of the time a fly spends in each subzone, normalised by the theoretical maximum value. The approach index was therefore defined as:

Approach index =

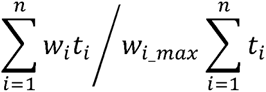

With:

*t*_*i*_: Time spent by the fly in subzone *i*.

*w*_*i*_: Weight assigned to subzone *i*, from 0.1 to 1.0 in ES zone (leftmost and rightmost subzones respectively), and 1.1 (in reward zone, termed *w*_*i*_*max*_)

*n*: The number of subzones (10 subzones in the ES zone plus the reward zone).

This formula therefore gives an approach index between 0 and 1, that gets higher as a fly goes further towards the reward zone. For example, a fly that rarely enters the ES zone and immediately goes back to the entry zone has an AI close to 0; a fly that spends more time in the middle of the ES zone but still doesn’t reach the reward zone will get a higher AI; and a fly that reaches the reward zone and spends a lot of time there will be attributed a high AI approaching 1. This way, if two flies reach the same point of the ES zone, the one that spent more time closer to the reward zone will therefore get a higher AI. Similarly, between two flies that both reached the reward zone, the one that spent the most time there will be attributed a higher AI. Thus, this formula allows for a robust measure of this complex avoidance-approach behaviour.

### CRTC::GFP immunohistochemistry

Five to ten days old male flies, either 24 hours starved or fed, expressing the UAS-CRTC: GFP construct were cold anaesthetised in a vial on ice and dissected in ice-cold 1x PBS. Cold anaesthesia and dissection were performed on a fly-by-fly basis to reduce time differences between anaesthesia and fixation. Dissected flies were fixed in 4% paraformaldehyde solution at room temperature for a period of 20 minutes. The fixed brains were then washed 4 times for 30 minutes each in 0.5% PBST (1x Phosphate-Buffered Saline, 0.5% Tween) before blocking for one hour in 5% normal goat serum. Following blocking, brains were then incubated with primary antibodies on a shaker for 3 days at 4°C anti-GFP chicken 1:1000 (Abcam, #13970) anti-DsRed rabbit 1:1000 (Takara, #632496). After incubation with primary antibodies brains were then washed four times in PBST for 30 minutes at room temperature. The brains were then incubated with secondary antibodies overnight at 4°C on a shaker Alexa Fluor 488 goat anti-chicken IgG 1:500 (ThermoFisher Scientific, #A28175), Alexa Fluor 546 goat anti-rabbit 1:500 (ThermoFisher Scientific, #A11035). After four 30-minute washes in PBST brains were mounted in Vectashield (Vector Labs) on a glass slide and covered with a coverslip. All brains were imaged on Zeiss LSM 900 with AiryScan2 module.

Calculation of the nuclear localisation index (NLI) was done as follows: (mean nuclear GFP signal) – (mean cytoplasmic GFP signal) / (mean nuclear GFP signal + mean cytoplasmic GFP signal).

### Two-photon functional imaging (Optogenetic stimulation and shock responses)

Three to six days old males expressing the calcium indicator GCaMP (details for genotypes on respective figures and legends) were dissected as in (*68*) and placed under a two-photon microscope (Sutter MOM customized by Rapp Optoelectronics). Fluorescence was generated by a Ti-Sapphire laser centered on 920□nm (Chameleon Vision S). Images with a pixel size of 0.3□×□0.3□μm were acquired with a Nikon 16x, 0.8 NA water-immersion objective, controlled by the ScanImage 2023.1.0 software (Vidrio/MBF Bioscience). Fast recordings were taken at a speed of 10□Hz with a resonant scan head (Sutter RGG). Analysis was performed using NOSA software v1.1.16 (neuro-optical signal analysis) (*69*), excel and Graphpad Prism. Regions of interest (ROIs) were manually drawn for analysis. Data was processed using a Savitzky–Golay filter. Mean intensity values were calculated as Δ*F*/*F*_0_, where *F*_0_ was defined as the mean *F* from baseline activity (first 10 s in Fig. 4F-H, Fig. 4G-K and Fig. 5C-E). The first 5 s (Fig. 4J-K) and 10 s (Fig. 5C-E) following imaging onset were discarded to eliminate motion and laser onset artefacts.

### Electric shock delivery under the two-photon microscope

The electric shock was delivered as 12 pulses of 75□V, each lasting 1.2 seconds, followed by a 3.8-second interval before the subsequent pulse. The electric shock grid was placed below the fly and controlled via a three-axis micromanipulator (Mini Manipulatorbloc, Luigs and Neumann). For calcium imaging, flies were head-fixed but positioned such that all legs contacted the electric grid, ensuring reliable shock delivery while still allowing leg movements. The fly was monitored via a monochrome USB3 camera (Blackfly S BFS-U3-13Y3M-C, FLIR) equipped with a megapixel varifocal lens (8.5–50 mm, f/1.6; A6Z8516CS-MP, Computer) and a near-infrared bandpass filter (BP735-40.5, MidOpt) during the recording. Illumination was provided from below by a high-power infrared LED (780 nm, 800 mW minimum output; M780LP1, Thorlabs), driven by a T-Cube LED driver (LEDD1B, Thorlabs). Calcium signals in FB2B neurons were recorded for a whole session of 180 s: 60 s before, 60□s during and 60□s immediately after the electric shock exposure. For experiments involving food delivery, food was presented to the fly in proximity during the whole imaging session without giving direct access (Fig. 4-5). Conditions under the microscope were set to > 20°C and > 40% humidity.

### Application of leucokinin or scrambled leucokinin peptide

Leucokinin (NSVVLGKKQRFHSWG-amide) and a scrambled control peptide (GNWSSVHVFLRGQKK-amide) were synthesized by GenScript. Peptides were diluted to 100 µM in sugar-free HL3 solution. Adult *Drosophila* were prepared by opening the head capsule to expose the brain. Peptide solution was applied directly onto the exposed brain and incubated for 5 min.

Calcium responses to food stimulation were recorded for 60 s before peptide application (pre-response) and for 60 s after peptide incubation (post-response).

### Optogenetic stimulation during vivo calcium imaging

Experiments were conducted at the Sutter MOM 2-Photon microscope with specifics mentioned above using an integrated 1-Photon UGA-42 Geo Module combined with a Laser Combiner LDI at 570 nm and 10% intensity. Calcium activity was recorded for 90 s expressing GCaMP6f or GCaMP7b in FB2B neurons while activating CsChrimson in PV5K1 neurons for 10 s at 15 s and at 45 s. The stimulation window was centered at FB2B presynapses where PV5K1 and FB2B neurons have contacts. Calcium signals during the stimulation windows were removed for clarity.

### Data visualisation

All data was visualised using GraphPad Prism 8.0.2 Boxplots show the median (centre line) and interquartile range (25th–75th percentiles); whiskers extend to the most extreme values within 1.5 × IQR (Tukey’s method). Calcium imaging traces over time are shown as mean ΔF/F_0_ (%; solid lines) with SEM (shaded area). To remove slow baseline drift in Fig. 5 caused by the 3-min continuous recording, a running 5th-percentile filter (50-s window) was subtracted from the ΔF/F, and the resulting baseline was zero-centered using the mean value during the first 50 s (pre-shock period). The Area Under the Curve (AUC) was calculated for the 60-second electric shock interval to quantify the total calcium response magnitude. AUC was determined using the baseline set at Y = 0.

### Statistical Analysis

Statistical analysis was performed using GraphPad Prism 10.6.1 and 8.0.2. The Shapiro-Wilk normality test was used to determine the normality of the data. The alpha value for all statistical tests was set at 0.05. After assessment of normality, paired t-tests or paired Wilcoxon signed-rank tests were used as appropriate. For comparisons involving more than two conditions, One-way ANOVA test followed by Tukey’s HSD post hoc was applied for the normally distributed data, and Kruskal–Wallis tests followed by Dunn’s post hoc was applied for the data not following normal distribution. Statistical analyses were performed on mean ΔF/F_0_ (%) values calculated for individual flies within the specific time windows indicated in the figures and/or Methods. Significant differences are indicated by asterisks. For each experiment, all genotypes were tested simultaneously, in random order, and at random times during the day to minimize potential effects of circadian phase and trial order. Outliers were identified using the ROUT method in GraphPad Prism (Q = 0.5%) and excluded prior to statistical analysis. See Suppl. Table 2 for details on statistics.

Behavioural choice data shown in Suppl. Fig. 1 were analysed using a binomial generalized linear model (GLM) with a logit link function. For each combination of shock voltage, feeding state, and availability food cues, flies were tested individually and outcomes were aggregated as the number of flies reaching the reward zone out of the total number tested. Shock voltage was treated as a continuous predictor, while feeding state (fed vs starved) and availability of food cues (food cues vs no food cues) were included as categorical factors. The full model included all main effects and interaction terms and was fit using maximum likelihood estimation. Statistical significance of model coefficients was assessed using Wald tests with heteroskedasticity-consistent (HC0) robust standard errors. The contribution of interaction terms was evaluated using likelihood-ratio tests comparing the full model to reduced models, including removal of the voltage × state × context term and removal of the state × context term while retaining voltage × state and voltage × context.

## Acknowledgements

We thank Isaac Cervantes Sandoval for insightful comments and critical assessment of the manuscript; Suewei Lin for helpful discussions; Andrew Lin, Daisuke Hattori, Emmanuel Perisse, and David Owald for sharing fly stocks; and members of the Rezaval lab, in particular Olga Procenko, as well as the Scheunemann lab, for valuable comments on the manuscript. We thank Dr Dean Kavanagh (Intravital Imaging Suite, RRID:SCR_027189), Technology Hub Facilities, College of Medicine and Health, University of Birmingham, for valuable technical assistance and input during this work, and Tzer Chyn Lim (BALM Imaging Facility, University of Birmingham) for assistance with light microscopy.

## Author Contributions

D.B. and C.R. conceived and designed the study. D.B. performed the behavioural experiments and functional imaging experiments using LK application and food responses. S.D., L.S. performed functional imaging experiments including shock responses and optogenetic stimulation. D.B. performed the fly genetics.□M.B. developed the approach index code. D.B and S.D. and L.S. analysed the data. D.B. and C.R. wrote the manuscript with input from M.B., S.D., L.S. C.R. provided supervision to D.B. and M.B., as well as overall supervision. C.R., L.S. provided resources. C.R. acquired funding.

## Data and materials availability

All data and code needed to evaluate and reproduce the results in the paper are present in the paper and/or the Supplementary Materials. Codes are available at https://github.com/lczl64/Bodas-et-al

## Supplementary Materials

**Supplementary Figure 1: Starvation and availability of food cues modulate reward approach across shock intensities** Reward-approach probability as a function of electric shock intensity. Points show observed probabilities (±95% binomial confidence intervals); lines show predictions from a binomial generalized linear model with voltage as a continuous variable. A significant voltage × feeding state × availability of food cues interaction (likelihood-ratio test, χ^2□^ = 7.63, p = 0.0057) demonstrates that the impact of aversive intensity on approach behaviour is contingent on motivational context. Specifically, starvation selectively alters the voltage–response relationship when food cues are available, whereas avoidance dynamics remain similar across all other conditions. A significant feeding state × availability of food cues interaction across voltages (χ^2□^ = 16.85, p= 4.0 × 10□□) further indicates that hunger modulates approach behaviour only in a reward-relevant context.

**Supplementary Figure 2. Silencing FB2B neurons does not alter basal walking velocity (A)** Expression pattern of the FB2B-split GAL4 driver labelling vFB neurons. **(B–D)** Mean locomotor velocity measured during the behavioural assay in FB2B-split GAL4 > UAS–TNT males and their respective genetic controls under fed conditions with food cues (**B**), fed conditions without food cues (**C**), and 24-h starvation without food cues (**D**). Silencing vFB neurons does not significantly affect walking speed across feeding states or food conditions (n = 27–29 flies per condition). Box plots indicate the median and interquartile range; whiskers represent the minimum and maximum values; points represent individual flies. Statistical significance was assessed using Kruskal–Wallis tests for panels B–D (ns, not significant).

**Supplementary Figure 3. Additional LK and LHLK characterisations (A-C)** Manipulation of LK neurons does not alter basal walking velocity. **(A)** Expression pattern of LK-GAL4 in the adult brain. **(B–C)** Mean locomotor velocity of LK-GAL4 males expressing UAS–Kir2.1 (**B**, 24-h starved) or UAS–dTRPA1 (**C**, fed) and their respective genetic controls, measured in the presence of food cues. Neither inhibition nor activation of LK neurons significantly alters locomotor velocity. Box plots indicate the median and interquartile range; whiskers represent the minimum and maximum values; points represent individual flies. (n = 30-33 flies per condition). Statistical significance was assessed using Kruskal–Wallis tests (ns, not significant). **(D)** Approach indices in fed LK-GAL4 > UAS–dTRPA1 males and their respective genetic controls tested at 24 °C during 75V electric shock with food cues (n=30-33 flies per condition). Statistical significance was assessed using Kruskal–Wallis tests (ns, not significant). **(E–F)** Starvation does not increase activity in SELK neurons, assessed using the transcriptional reporter CRTC::GFP. Representative images (**E**) and quantification of nuclear localisation (**F**) are shown (n = 7 flies per condition). Statistical significance was assessed using Mann–Whitney test (ns, not significant). Box plots indicate the median and interquartile range; whiskers represent the minimum and maximum values; points represent individual flies.

**Supplementary Figure 4. Additional characterisations of FB2B neurons and LHON neurons (A-D)** Manipulation of LHONs does not alter basal walking velocity. **(A)** Expression pattern of LH1478-GAL4 targeting PV5K1 neurons in the brain. **(B-D)** Mean locomotor velocity of LH1478-GAL4 males expressing UAS–LKR RNAi (B), UAS–TNT (C), or UAS–ChR (D) and their respective genetic controls under fed or starved conditions in the presence of food cues (n = 27–30 flies per condition). Statistical significance was assessed using Kruskal–Wallis tests (ns, not significant). **(E-F)** LK signalling in FB2B neurons is not required for food approach during ES. **(E)** Expression pattern of FB2B-GAL4 targeting FB2B neurons in the ventral fan-shaped body (vFB). **(F)** Downregulation of leucokinin receptors in FB2B neurons does not alter food approach in starved flies. Approach indices were measured in 24-h-starved FB2B-split GAL4 > UAS–LKR RNAi males and their respective genetic controls tested in the presence of food cues (n = 19-20 flies per condition). Box plots in B-D, F indicate the median and interquartile range; whiskers represent the minimum and maximum values; points represent individual flies. Statistical significance was assessed using a Kruskal–Wallis test (ns, not significant). **(G)** Application of 100 µM LK peptide alone does not change GCaMP signal in PV5K1 neurons. Δ*F*/*F*_0_ traces of LH1478-GAL4>UAS-GCaMP6f signals in fed flies, before and after application of 100 µM LK peptide (n = 7 flies per condition). Statistical significance was assessed using paired Wilcoxon signed-rank test (ns, not significant).

**Supplementary Video 1.** Representative example of the approach–avoidance assay showing fed male wild-type flies encountering an electrified shock zone in the presence of food. Unlike starved flies, fed flies consistently avoid the aversive zone and fail to reach the food reward.

**Supplementary Video 2.** Representative example of a 24-h-starved male wild-type flies traversing an electrified shock zone to reach a food reward. In the presence of food, hungry flies suppress nociceptive avoidance and cross the aversive zone to access the reward.

**Supplementary Table 1.** List of strains and genotypes.

**Supplementary Table 2.** Statistics for behavioural, CRTC::GFP and live calcium imaging data (main and extended figures).

