## Supplementary figures and images for "A state-dependent neural circuit resolves approach–avoidance conflicts"

### Suppl. Figure 1

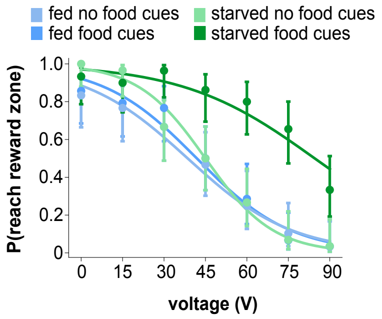


Suppl. Figure 1

### Suppl. Figure 2

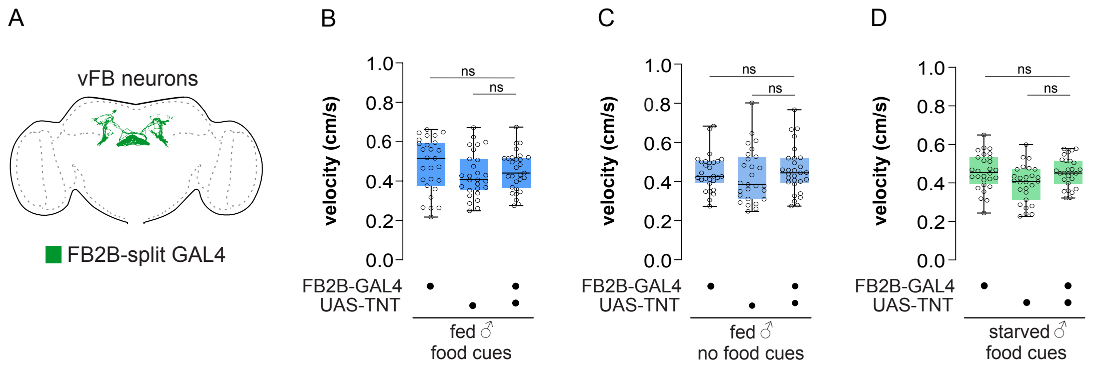


Suppl. Figure 2

### Suppl. Figure 3

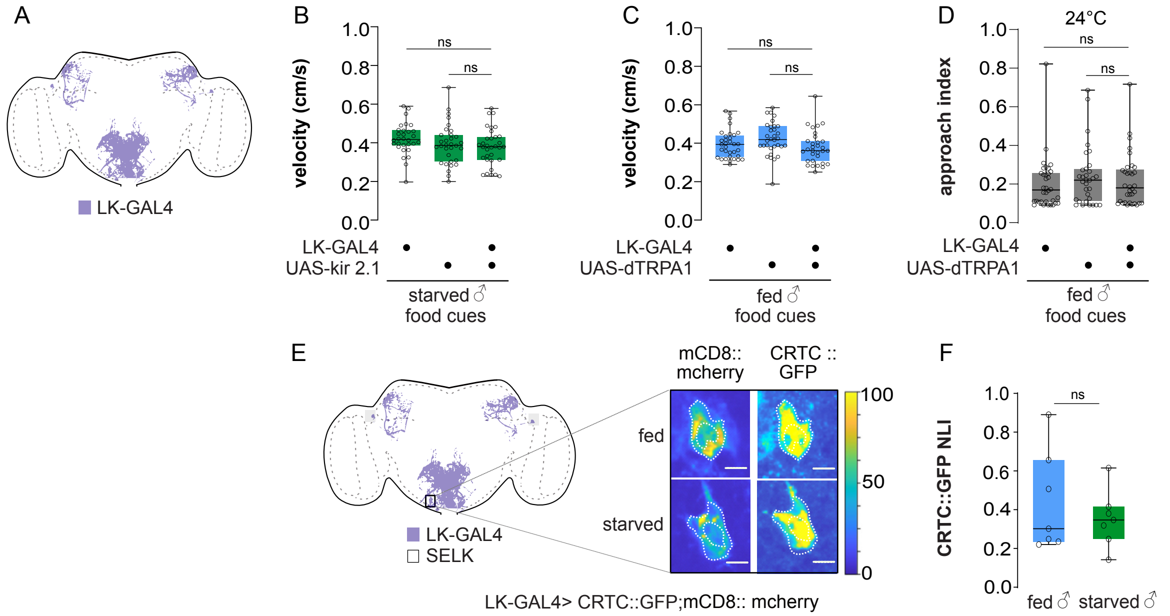


Suppl. Figure 3

### Suppl. Figure 4

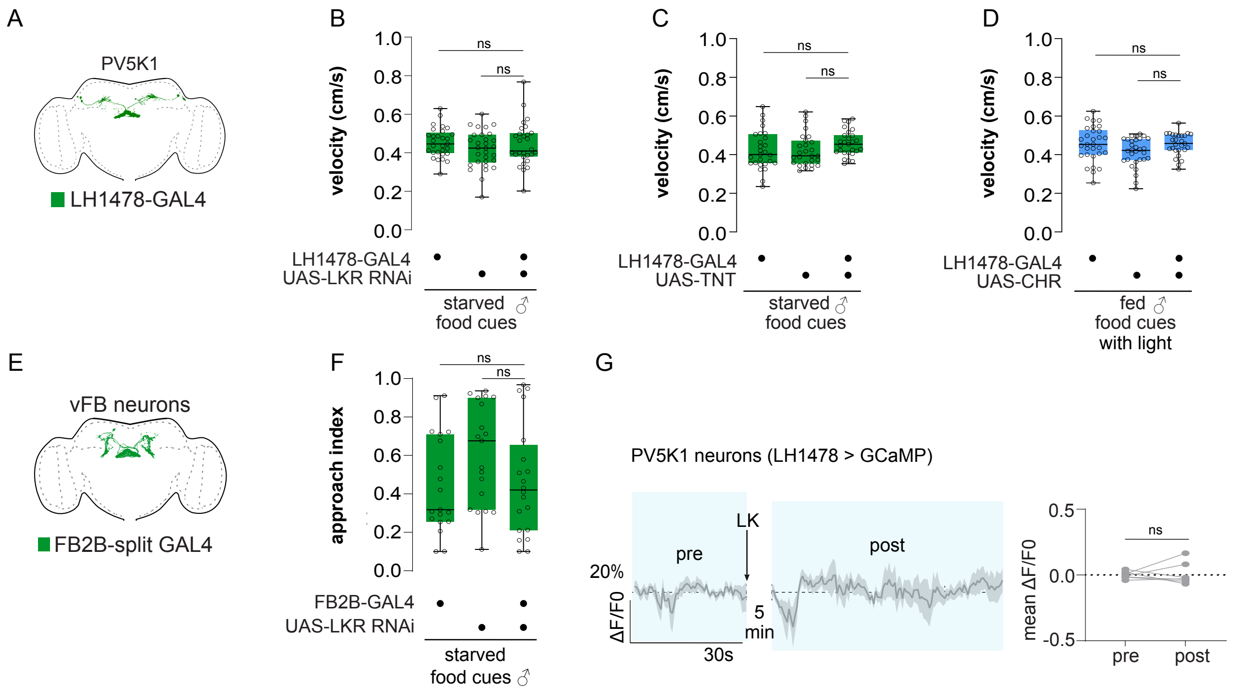
